# A prior-based approach for hypothesis comparison and its utility to discern among temporal scenarios of divergence

**DOI:** 10.1101/302539

**Authors:** Eugenia Zarza, Robert B. O’Hara, Annette Klussmann-Kolb, Markus Pfenninger

## Abstract

One of the major problems in evolutionary biology is to elucidate the relationships between historical events and the tempo and mode of lineage divergence. The development of relaxed molecular clock models and the increasing availability of DNA sequences resulted in more accurate estimations of taxa divergence times. However, finding the link between competing historical events and divergence is still challenging. Here we investigate assigning constrained-age priors to nodes of interest in a time-calibrated phylogeny as a means of hypothesis comparison. These priors are equivalent to historic scenarios for lineage origin. The hypothesis that best explains the data can be selected by comparing the likelihood values of the competing hypotheses, modelled with different priors. A simulation approach was taken to evaluate the performance of the prior-based method and to compare it with an unconstrained approach. We explored the effect of DNA sequence length and the temporal placement and span of competing hypotheses (i.e. historic scenarios) on selection of the correct hypothesis and the strength of the inference. Competing hypotheses were compared applying a posterior simulation analogue of the Akaike Information Criterion and Bayes factors (obtained after calculation of the marginal likelihood with three estimators: Harmonic Mean, Stepping Stone and Path Sampling). We illustrate the potential application of the prior-based method on an empirical data set to compare competing geological hypotheses explaining the biogeographic patterns in *Pleurodeles* newts. The correct hypothesis was selected on average 89% times. The best performance was observed with DNA sequence length of 3500-10000 bp. The prior-based method is most reliable when the hypotheses compared are not temporally too close. The strongest inferences were obtained when using the Stepping Stone and Path Sampling estimators. The prior-based approach proved effective in discriminating between competing hypotheses when used on empirical data. The unconstrained analyses performed well but it probably requires additional computational effort. Researchers applying this approach should rely only on inferences with moderate to strong support. The prior-based approach could be applied on biogeographical and phylogeographical studies where robust methods for historical inferences are still lacking.

## Introduction

One of the major problems in evolutionary biology is to elucidate the relationships between historical events and the tempo and mode of lineage divergence and, ultimately, biological diversification. The development of methods to estimate substitution rates with relaxed molecular clock models and the increasing availability of DNA sequences has led to better estimates of species and higher taxa divergence times (Battistuzzi et al., 2010). However, finding the link between historical events, such as past geological and climatic changes, and divergence is still challenging. As phylogeography – and other evolutionary biology disciplines - move away from narrative and traditional null-hypothesis methods towards multiple hypothesis comparison approaches (Johnson & Omland, 2004; Dépraz et al., 2008; Bloomquist, Lemey & Suchard, 2010; Carstens et al., 2013), it is necessary to investigate if a hypothesis comparative framework can also be applied at deeper evolutionary times.

Hypothesis comparison offers a means to draw inferences from a set of multiple competing hypotheses and to estimate the degree of confidence that can be placed on each of them (Dépraz et al., 2008; Johnson & Crandall, 2009). Competing hypotheses should be thoroughly thought through and formulated as a first step in the research process (Anderson, 2008). After data collection and analyses, the competing hypotheses can be compared and ranked to select which of them best explains the data. This can be accomplished using the Bayes factor (BF), the ratio of the marginal likelihood of the data from two models, i.e. the posterior probability of one model to that of another model, divided by the ratio of the prior probabilities, thus BF measures the change in support for one model versus another given the data (Jeffreys, 1935; Kass & Raftery, 1995; Suchard, Weiss & Sinsheimer, 2001, 2003).

Here we propose a hypothesis comparison approach to evaluate the influence of historic events in lineage divergence. Our main aim is to explore if assigning constrained-age priors to nodes of interest in a time-calibrated phylogeny would serve as a means for hypothesis comparison. These priors would be equivalent to scenarios for lineage divergence under certain competing hypotheses. When comparing the likelihood values of such hypotheses, modelled under different priors, we would be able to select the hypothesis that best explains the data and assign a level of confidence to evolutionary inferences. This hypothesis comparison approach has been employed a few times to empirical data with success to discern among competing temporal biogeographical scenarios in crabs (Klaus et al., 2010; Jesse et al., 2011), and land snails (Pfenninger et al., 2010). However, the efficiency, accuracy and range of validity of the approach have as yet not been rigorously tested in a systematic manner.

### METHODS FOR MODEL SELECTION: BAYES FACTORS AND AKAIKE’S INFORMATION CRITERION

Bayes factors allow for hypothesis ranking and evaluation of the relative merits of the competing hypotheses (Jeffreys, 1935; Kass & Raftery, 1995; Baele et al., 2012), placing BF at the core of Bayesian theory of hypothesis (Robert & Wraith, 2009). When using BF, the model, or in this case hypothesis, with the greatest marginal likelihood (for simplicity MLL) is generally preferred. The marginal likelihood is a weighted average of the likelihood, where the weights come from the prior (Xie *et al*. 2011). In a phylogenetic context where the parameter space is very large, calculating MLL, requires integrating over all possible solutions and is not analytically feasible.

Until recently, importance sampling approaches were used to calculate the harmonic mean estimator (HM) of MLL (Newton & Raftery, 1994), despite the short-comings of the approach being outlined in the original paper. HM only needs simulations from the posterior distributions and can be easily calculated from an MCMC sample. Consequently, it has been widely used in phylogenetics (e.g. MrBayes and implemented in BEAST). However, HM is not stable and can have infinite variance giving unreliable results for model selection (Lartillot & Philippe, 2006; Xie et al., 2011). Recent developments aim to improve the exploration of the relevant model space via guided transitions across a sequence of intermediate distributions connecting their prior and posterior extremes (Cameron & Pettitt, 2013). Among these methods are thermodynamic integration (Lartillot & Philippe, 2006), also known as path sampling (PS; Ogata, 1989; Gelman & Meng, 1998) and the Stepping Stone method (SS; Xie et al., 2011). Both methods have been implemented in BEAST latest version (from version 1.7.0; Drummond et al., 2012; Baele et al., 2012), together with a posterior simulation analogue of the Akaike’s information criterion through MCMC (AICM; Raftery et al., 2007; Baele et al., 2012), forming a useful set of tools for model selection in phylogenetics.

Here, using a simulation approach, we evaluate the plausibility of using prior information to compare hypotheses on divergence times between lineages. We apply several model selection techniques (AICM, HM, SS and PS) and evaluate their performance for prior-based hypothesis comparison under several conditions. We varied data amount, relative temporal placement, span and absolute tree location of hypotheses (age priors), but kept the evolutionary and relaxed clock models constant. Using a reduced set of simulations, we compared the prior-based approach with a simpler approach to select among competing scenarios. This consists in executing one analysis to compute the proportion of sampled MCMC steps that fall within date intervals compatible with competing historical events or scenarios. In this way, for each scenario, it would be possible to estimate the posterior probability that a given divergence occurred at the same time as the historical event. This would enable comparing several competing hypotheses through BF. This approach does not require applying constraints on the age prior distribution of the nodes of interest and we refer to it as the “unconstrained” analysis (Uc).

We illustrate the potential application of the prior-based hypothesis approach on an empirical data set to compare competing geological hypotheses explaining the biogeographic patterns of *Pleurodeles* newts in Iberia and Northern Africa (Zhang et al., 2008).

## Materials and Methods

A simulation study was carried out to evaluate the performance of the prior-based hypothesis comparison approach, to investigate what factors could lead to a reliable selection of the correct hypothesis or scenario and to compare it with Uc. Three sets of simulations were generated. The main difference among them is the age of the simulated correct hypothesis: “deep” correct hypothesis (DCH, 4.2-4.7. Ma), “intermediate” correct hypothesis (ICH, 2.7-3.2 Ma) or “shallow” correct hypothesis (SCH, 1.2-1.7 Ma). The general simulation procedure included several stages. The first step was to simulate trees with 25 taxa with BEAST v1.6.1 (Drummond et al., 2006) in which five nodes were age-constrained to the same time interval (e.g. 2.7-3.2 Ma, the onset of the Northern Hemisphere Glaciation NHG). We constrained this high number of nodes to facilitate divergence time estimation and reflect a scenario where many nodes in the tree were affected by a very significant event. Nodes were defined as two-taxa set, with a uniform-age prior reflecting the age of the event. Trees were built following a birth-death tree prior and a normal prior with 15 Ma mean for the root age, without sequence data. To test performance under a variety of tree shapes, trees were sampled at a frequency resulting in up to 100 final trees from which 20 were randomly selected, using the random.org number generator. All the nodes in the trees were resolved and the topologies and position of the nodes in the tree are shown in File S1. Input files to generate the simulated trees are in File S2.

The topologies of the selected trees were used to simulate DNA sequence data with Seq-Gen (Rambaut & Grassly, 1997). Five partitioned DNA-sequences datasets of 3500, 10000 and 20000 bp were simulated for each topology under the Jukes-Cantor substitution model. To reflect partition rate heterogeneity, we specified a relative rate of evolution for each partition. As required by SeqGen, the relative rates had a mean of 1.0, but without variation in the substitution rate among taxa. Number of partitions per data set is shown in Table S1. Each data set was used to generate input files for BEAST. The age priors for the nodes of interest in these input files reflect the “correct” hypothesis (i.e. DCH, ICH or SCH), which has the same age priors as those used to generate the simulated topology. The sequence data was also used to create input files for BEAST with age priors reflecting the competing hypotheses described in the following sections (supplementary Table S2, Fig. 1 and Fig. 2). Input files reflecting the correct and competing hypotheses had additional time calibrations on one or two nodes and the tree root. These nodes’ age prior follows a normal distribution, whose mean corresponds to the age of that node in the initial simulated topology. In a similar way, input files were created for the unconstrained analyses and are included in File S2 to facilitate analysis replication.

**Figure 1.**
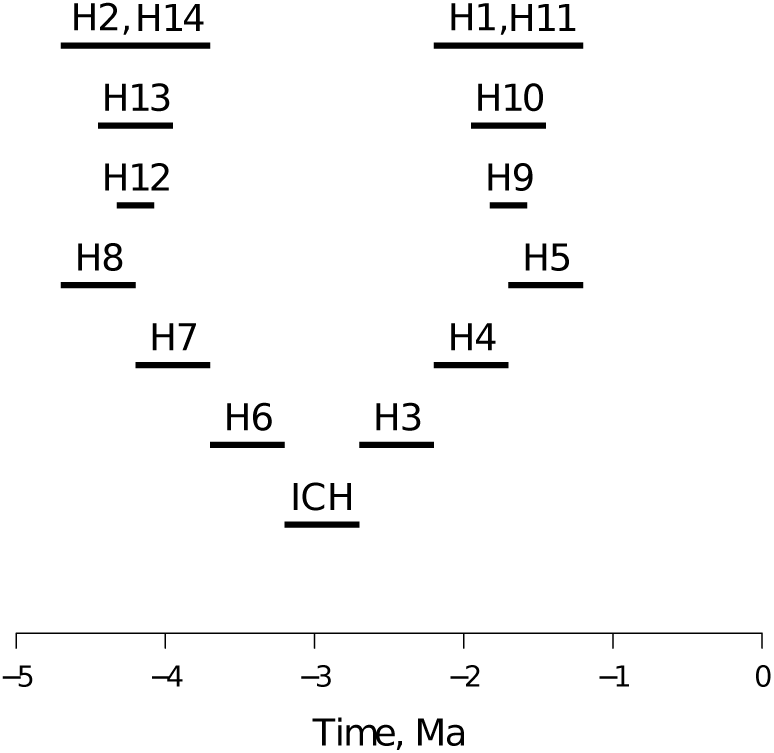
Competing hypotheses. Lines represent the temporal location and span of competing hypotheses. ICH= correct hypothesis; H1-H14 competing hypotheses (see also Table S2); Ma=million years ago.

**Figure 2.**
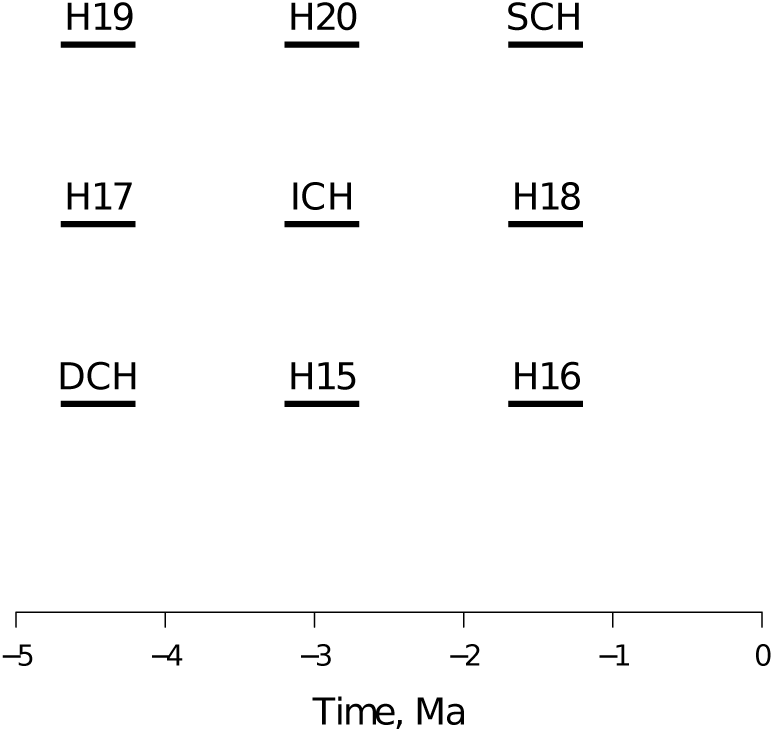
Variations in temporal depth of correct hypotheses. Lines represent the temporal location of the deep (DCH), intermediate (ICH) and shallow (SCH) correct hypotheses, with their respective competing hypotheses shown in the same row.

The analyses were run under an uncorrelated relaxed molecular clock (UMC). Although the sequences were generated without variation in the substitution rate among taxa, it has been suggested that UMC reliably estimates parameters even when the data follows a strict molecular clock, which is indeed a model comprised within the more complex UMC (Drummond et al 2006). Thus we do not consider that this will be detrimental for this study. As the substitution rate is unknown for most no-model organisms, we consider that it will be more informative to estimate this parameter from the data.

### EFFECT OF SEQUENCE LENGTH, HYPOTHESIS RELATIVE TEMPORAL POSITION AND HYPOTHESIS TEMPORAL SPAN

In this set of simulations, the correct hypothesis age was fixed to the intermediate time depth (i.e. ICH, 2.7-3.2) whereas sequence length and position and temporal span of competing hypotheses varied. Three historical scenarios were compared: 1) nodes split at the time of a geological/climatic event: the time of the NHG at 2.7-3.2 Ma, and is considered as the correct hypothesis ICH; 2) split occurred before the geological/climatic event; 3) split occurred after the geological/climatic event. These scenarios reflect the situation of a researcher who suspects that a climatic/geological event might have led to a node split in a phylogeny, but would like to know how much better (or worse) the hypothesis explains the data in comparison to the other scenarios.

To test sequence length effect of on hypothesis selection, we simulated data sets with 3500, 10000 and 20000 bp and compared ICH to competing scenarios where nodes split before or after ICH. To explore how temporally close the competing hypotheses and ICH can be to properly distinguish and select ICH, we used a more or less intermediate data set size (i.e. 10000 bp) and varied the temporal location of the competing hypotheses one or two intervals before and after ICH, thus competing hypotheses did not overlap. An interval is defined as equal to the temporal span of ICH: 0.5 million years (Myr). Another important factor to consider is the competing hypotheses duration, specifically is it valid to compare hypotheses with different widths of prior age distributions? To answer this question, we simulated competing hypotheses where the age priors of the nodes of interest were two times wider than, equal to, or half as wide as ICH, and were temporally located before and after ICH.

One hundred replicate input files were generated for each type of competing hypothesis (temporal or duration variation), following the general procedure above described. Input files were run in BEASTv1.7.1. MCMC length is shown in supplementary Table S1. The comparable competing hypotheses were run with the same number of iterations.

### EFFECT OF ABSOLUTE AGE OF CORRECT HYPOTHESIS

To test how correct hypothesis absolute age (i.e. temporal depth) affects hypothesis comparison and selection, we followed the general simulation procedure. Tree topologies were simulated where five nodes of interest were constrained with age priors reflecting DCH or SCH. For each situation, the 20 randomly chosen trees were used to generate DNA-sequence data sets of 10000 bp. DCH was compared to more recent competing hypotheses: 2.7-3.2 Ma and 1.2-1.7 Ma; whereas SCH was compared to older competing hypothesis: 2.7-3.2 Ma and 4.2-4.7 Ma. Performance with these two variations of correct hypotheses was compared to performance with ICH. We removed runs that did not converge to keep the run length equal among simulations with similar data set size.

### HYPOTHESIS SELECTION

The marginal likelihood under the different priors was estimated using the HM, PS and SS methods. The natural logs of the Bayes factors were calculated as ln(BF)=*H_i_*-*H_j_*, where *H_i_* and *H_j_* are the log natural of the competing hypothesis MLL, following the method first implemented in Tracer (Suchard, Weiss & Sinsheimer, 2003; Rambaut & Drummond, 2007) to calculate BF based on HM. The strength of evidence was evaluated according to the table provided by Kass and Raftery (1995) but without multiplying by 2 and without rounding up ln(BF) values). Thus, ln(BF)<1.10 means weak support for *H_i_* over *H_j_*, 1.10 < ln(BF) < 2.30 mean moderate support and a ln(BF) > 2.3 was considered as strong support (BF >10). Regarding selection with AICM, a Δ AICM >10 between the best ranked hypothesis and the other hypotheses suggests that the latter were very unlikely (Burnham & Anderson, 2002). These calculations where performed in BEASTv1.7.1 with the code of Baele *et al*. (2012). It is expected that the “correct” hypothesis will have higher MLL values than the others if our method is effective.

### UNCONSTRAINED ANALYSES

The frequency of the MCMC steps falling within each of the correct and competing hypotheses time intervals was calculated to estimate the posterior probability of each hypothesis. We calculated the prior probability of a hypothesis as its interval length divided by the total interval length (i.e. the time from its most recent calibrated ancestral node to the present). BFs were calculated as the ratio of posterior odds to prior odds between the correct hypothesis and a particular competing hypothesis. This was obtained for each of the five nodes, for each competing hypothesis only for the treatments comparing against ICH, and with data sets of 3500, 10000 and 20000 bp. To make Uc results comparable with the prior-based approach, we estimated inference strength with this scale: BF<1 false positive; 1< BF <3.01 weak; 3.01 < BF < 10 moderate; BF > 10 strong. The frequency of strong, moderate and weak BF per node was calculated. We calculated an average frequency of strong, moderate and weak BF for the five nodes, for each treatment.

### EMPIRICAL DATASET ANALYSIS: SALAMANDERS

In this section we apply the prior-based approach to compare hypotheses on the time of split between two species of newts and the influence of geological and climatic events. Zhang *et al* (2008) proposed a time-calibrated phylogeny of the family Salamandridae inferred from mitochondrial genomes (10755 bp). The data set comprises 41 taxa, including representatives of all recognized genera. The authors calibrated six nodes with fossil records and one using indirect geological evidence. Based on the results of Bayesian and penalized likelihood analyses the authors proposed a robust time-calibrated phylogeny and postulated several biogeographic hypotheses to account for the distribution patterns between taxa in Salamandridae. We re-analysed their data set to compare three previously suggested competing scenarios to explain the phylogeographic patterns observed in one of the clades, the ribbed newts (*Pleurodeles*), currently distributed in Iberia and Northern Africa (Frost, 2011). According to Veith *et al* (2004) and Zhang *et al*. (2008): 1) The split between *P*. *waltl* and *P*. *poireti* could be consistent with the Messinian salinity crisis (ca. 5.33 Ma); or 2) The Betic crisis ca. 14 Ma; or 3) the Betic crisis leading to the split between the north-western and south-eastern populations of *P*. *waltl*, rather than between the two *Pleurodeles* species, which would imply that the two species split around 35 Ma.

We used the BEAST input file of Zhang *et al*. (2008) keeping the original fossil calibration points but assigning proper priors to all parameters (Baele *et al*. 2013). In three independent analyses, age priors were added to reflect the competing scenarios. Analysis 1 included the original calibration points plus a normal age prior for the most recent common ancestor of *Pleurodeles* species (from now on referred to as Node P) with mean 5.33 Ma, reflecting scenario 1. In analysis 2, in addition to the original calibration points, a normal age prior was assigned to Node P with mean 14.0 Ma, reflecting scenario 2. In analysis 3, the original calibration points plus a normal age prior with mean 35 Ma, reflecting scenario 3, were included. To obtain adequate effective sample sizes of the parameters, five independent runs with 100 million MCMC iterations were executed in BEASTv1.7.1. After MCMC execution, samples of the prior and posterior were collected for later estimation of MLL with HM, PS and SS, following suggestions on the BEAST website (beast.bio.ed.ac.uk/Model_selection). Log files of the five independent runs were combined with LogCombiner of the BEAST package after removing 10% of the samples as burnin. The combined log files were used to calculate the AICM and estimate MLL using HM. PS/SS analyses were executed combining the samples of power posteriors collected at the end of each MCMC. Competing scenario MLLs were then calculated to select the one that best explains the data.

## Results

One hundred simulated replicate datasets were analysed per “treatment”. However, convergence of the MCMC runs for the alternative, the correct hypotheses or/and the unconstrained analyses was not always achieved and acceptable effective sample sizes were not obtained. In the prior-based approach, runs that failed to converge and their competing hypotheses (correct or alternative hypotheses)-even if these converged-were not taken into account to calculate the effectiveness of the method. An improvement of up to 5% in the frequency of success was observed when ignoring the runs lacking convergence in comparison to keeping all runs irrespective of convergence achievement. The PS and SS methods produced similar results under all the simulations strategies, thus only one graph is shown.

The unconstrained analyses consisted on executing one run to compare the frequency of MCMC steps falling within the intervals of several competing hypothesis. The Uc analyses were run for the same number of MCMC iterations as the prior-based approach. However with data sets of 3500, 10000 and 20000bp, 7%, 39% and 49% of the Uc runs did not reach convergence, respectively; whereas in average 0%, 8.7% and 18% of the respective competing hypothesis runs in the prior-based approach did not converge.

### EFFECT OF SEQUENCE LENGTH

Sequence length was increased from 3500 bp up to 20000 bp as shown in supplementary Table S1. With the prior-based approach, all the MCMC runs analysing 3500 bp data sets achieved convergence. Runs of the correct hypothesis and its competing hypotheses reached convergence 78% and 59% with 10000 bp and 20000 bp data sets, respectively. Increasing sequence length leads to an increase in the frequency of selecting the correct hypothesis as the best hypothesis with strong support when using AICM and HM (Fig. 3). However an improvement is not seen when calculating MLL with PS/SS with data sets larger than 10000 bp (Fig. 3). False positives frequency decreases with sequence length from 3500 bp to 10000 bp with all methods (HM: from 12.5 to 4.6 %; AICM: from 9.5% to 3.9%; SS/PP from 6.5% to 5.9%). Only with AICM can a reduction in false positives be seen with 20000 bp data sets (3.3%). Nevertheless, strong inferences frequency is always higher when using PS/SS than with HM, and AICM (Fig. 3). Uc shows better performance than HM and AICM with 10000 and 20000 bp data sets, but performs poorly with small data sets.

**Figure 3.**
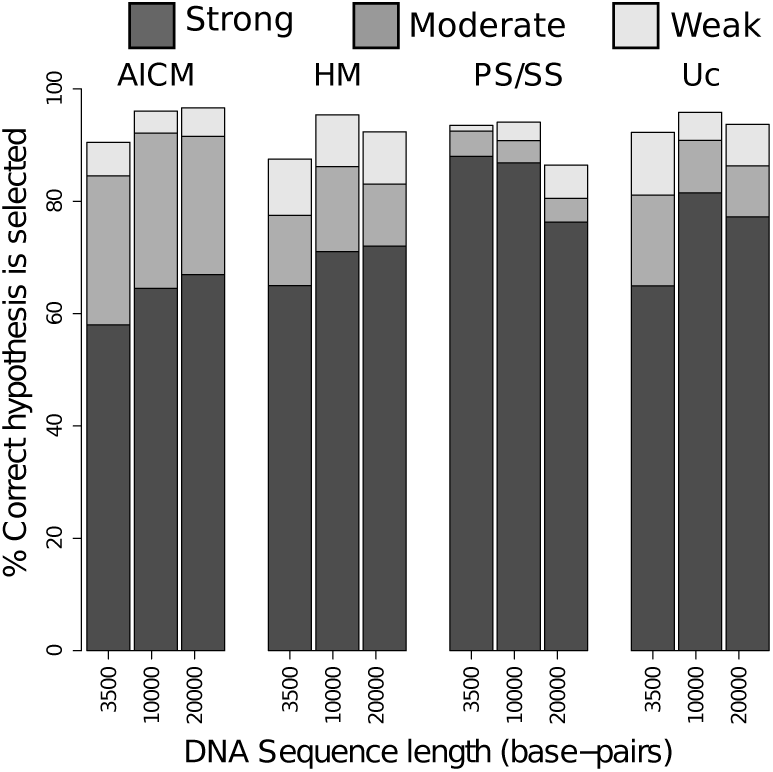
Effect of sequence length on selecting the correct hypothesis. Bars represent the average frequency of ranking ICH as the best hypothesis and strength of inference according to the method employed.

### EFFECT OF TEMPORAL SPAN OF COMPETING HYPOTHESES

Different sizes for the temporal constraint interval of the competing hypotheses were compared. Regarding the prior based approach, convergence was achieved by 78% of the correct hypothesis and its competing hypotheses MCMC runs. With the AICM calculation, the correct hypothesis was selected above 96% of the times, with no strong false positives. HM performs with a similar rate of success, however the correct hypothesis is selected with strong support more often than with AICM with only 0.65% of strong false positives. In both cases a better performance was obtained when the hypotheses span intervals of similar size or when the competing hypothesis has a narrower temporal range. PS/SS select the correct hypothesis strongly more frequently than the other two methods (Fig. 4). The correct hypothesis was strongly supported slightly more often (90%) when the competing hypotheses had narrower intervals than when the competing hypotheses had an interval as wide as the correct hypothesis (87%; Fig 4). Strong false positives were obtained at a frequency between 2.6 to 3.9%. It should be noted that the AICM does not estimate MLL and thus the results are not entirely comparable. Uc performed better than AICM and HM with all interval sizes and was slightly outperformed by PS/SS.

**Figure 4.**
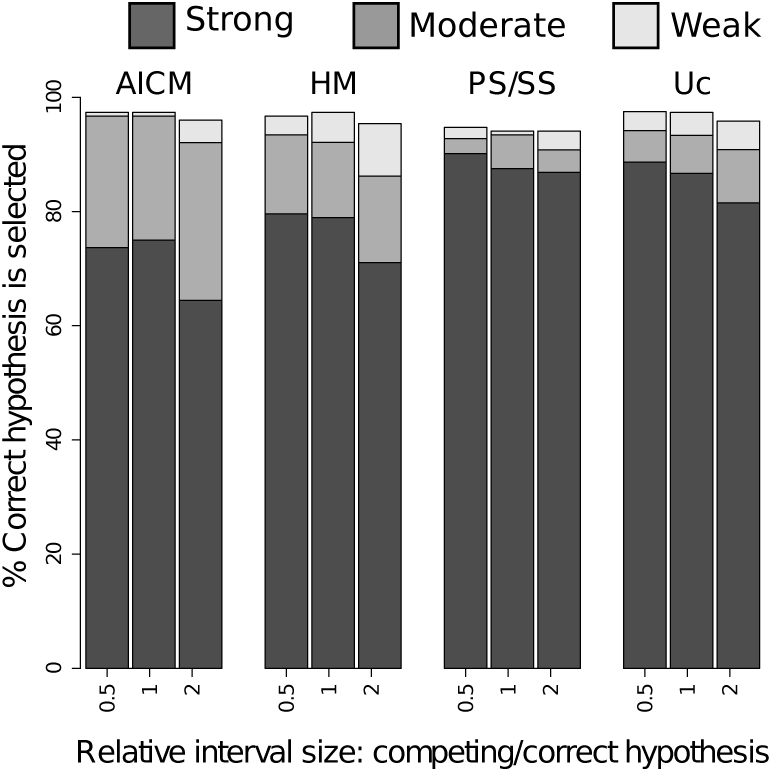
Effect of temporal span (relative interval size) of competing hypotheses on selecting the correct hypothesis. Bars represent the average frequency of ranking ICH as the best hypothesis and strength of inference according to the method employed. Interval=0.5 Million years (Myr).

### EFFECT OF RELATIVE TEMPORAL LOCATION OF HYPOTHESIS

Temporal location of competing hypotheses was also varied. In the prior-based approach, convergence was achieved in 76% of the correct hypothesis and its competing hypotheses runs. Our simulations suggest that the closer the competing hypothesis is to the correct hypothesis the less likely it will be to rank the correct hypothesis as the best hypothesis (Fig. 5). A trend towards increase in selection accuracy with increase in temporal distance between hypotheses was observed with all methods. BFs calculated with PS/SS select the correct hypothesis with strong support more often than HM when the hypotheses are the furthest apart (92% and 86% respectively). PS/SS produce stronger inferences than HM when the hypotheses are the closest, although the performance is poor (<50%). Selection of the correct hypothesis with AICM with moderate to strong support occurs above 78% of the times when hypotheses are the furthest apart. High frequency (19%) of false positives was observed when applying HM and hypotheses were very close together, but they are reduced when the hypotheses are further apart (1.9%). False positives frequency obtained with AICM is reduced from 18% to 2.6 % when hypotheses are the furthest away. PS/SS produce the highest frequency of false positives when the hypotheses are close (8.3%), but this is reduced when the hypotheses are temporally apart (0.64%). Similarly, with Uc it is difficult to select among closely located hypotheses.

**Figure 5.**
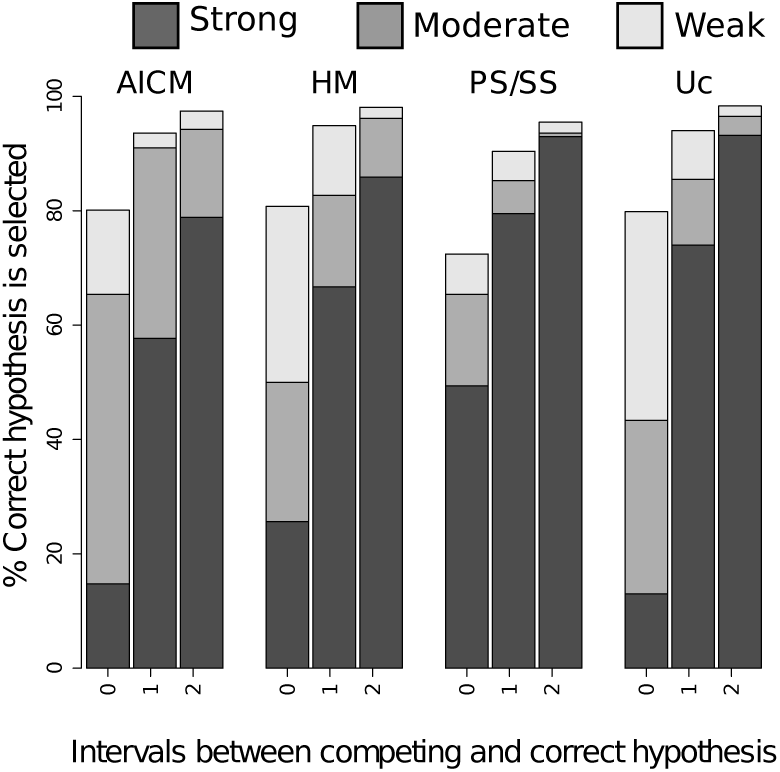
Effect of temporal location of competing hypothesis on selecting the correct. Bars represent the average frequency of ranking ICH as the best hypothesis and strength of inference according to the method employed Interval=0.5 Myr

### EFFECT OF ABSOLUTE AGE OF THE CORRECT HYPOTHESIS

To investigate the effect of the absolute age of correct hypothesis in the tree, two sets of simulations were carried out. The first simulated SCH and was compared with less recent hypotheses. In this case, 90% of the MCMC runs achieved convergence. All four methods selected SCH as the correct hypothesis 100% of the times with strong support (Fig. 6 A). Strong false positives occurred only in one case when using PS/SS and HM. When the correct hypothesis was ICH, AICM and HM tended to perform better when the competing hypothesis is deeper than ICH, with a higher frequency of strong inferences. PS/SS led to stronger inferences over more recent hypotheses (Fig. 6 B). No strong false positives were obtained except for one case when using PS/SS. Only 15% of the runs reached convergence in simulations with DCH. Among these runs, PS/SS performed better than the other two methods selecting the DCH above 93% of the times with strong support, followed by AICM and HM (Fig. 6 C).

**Figure 6.**
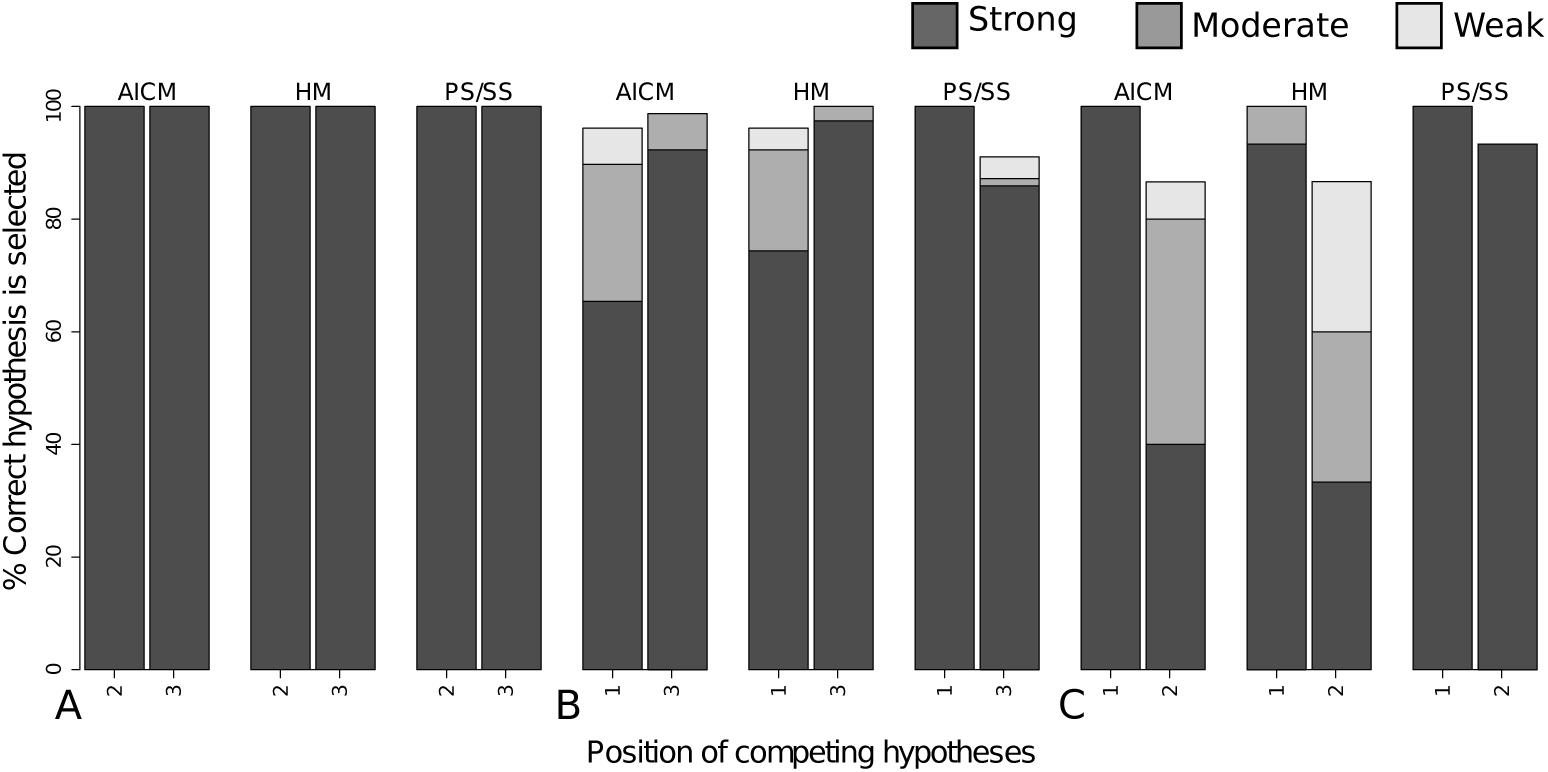
Effect of absolute age (temporal depth) of the correct hypothesis. A) Frequency of selecting the shallow age correct hypothesis (temporal position 1) over deeper competing hypotheses (temporal position 2 and 3). B) Frequency of selecting the intermediate age correct hypothesis (temporal position 2) over a shallower (temporal position 1) and a deeper competing hypothesis (temporal position 3). C) Frequency of selecting the deep age correct hypothesis (temporal position 3) over shallower competing hypotheses (temporal position 1

### HYPOTHESIS COMPARISON USING EMPIRICAL DATA

After combining the MCMC outputs, effective sample sizes above 100 were obtained for all parameters. The three independent competing analyses resulted in the same topology obtained by Zhang *et al*. (2008). Evidence is stronger for Scenario 2 when BFs are estimated based on MLL calculated with PS, SS and HM methods (Table 1). However, AICM ranks Scenario 3 as the best hypothesis. The Bayes factors calculated with PS and SS estimates are larger than those obtained with HM. Δ AICM moderately supports Scenario 3 over the other competing hypotheses. The results from PS, SS and HM are in agreement with results previously obtained with molecular and fossil evidence, suggesting that the split between these species of ribbed newts is associated with the Betic crisis (Zhang et al., 2008).

**Table 1.** Comparison between hypothesised scenarios for the time of split between *Pleurodeles*, using Bayes Factors calculated based on HM, PS, SS Marginal Likelihood estimates and Δ AICM. A value 0 indicates the best ranked hypothesis.

| Scenario | Bayes<br>Factors<br>(PS) | Bayes<br>Factors<br>(SS) | Bayes<br>Factors<br>(HM) | Delta<br>AICM |
| --- | --- | --- | --- | --- |
| 1 | -9.96 | -11.06 | -0.12 | -1.50 |
| 2 | **0** | **0** | **0** | -2.72 |
| 3 | -13.8 | -15.53 | -0.31 | **0** |

## Discussion

We evaluated the performance of a hypothesis comparison approach that uses prior information to define competing scenarios of lineage divergence, in which divergence is associated with historic events like climate or geological change. After calculation of their marginal likelihood or AICM, it is possible to rank scenarios and select the one that best explains the data. Our simulation study suggests that under reasonable circumstances, this approach could constitute a reliable tool to compare temporal scenarios: the correct hypothesis is ranked as the best hypothesis over 80% of the time under almost all simulation strategies. However, inference strength varies depending on the method employed to calculate BF or if AICM is used. Most of the times HM ranks the correct hypothesis as the best hypothesis but the BFs are so low that it is difficult to place any confidence in the selection. Generally, PS and SS estimates of MLL differ more strongly between competing hypotheses than HM. We observed that these methods could also lead to few false positives with strong or moderate support. This may, in part, be because the data genuinely support the wrong hypothesis by chance (e.g Kuparinen et al., 2007).

Discerning between competing hypotheses is particularly challenging when the hypotheses are located close to each other in time. Interestingly, it was consistently difficult to reach convergence when the node of the correct hypothesis was located deeper in the tree (DCH), especially for runs where the alternative hypotheses were the furthest away from the correct hypothesis. The accuracy and strength of ranking the correct hypothesis as the best hypothesis increase slightly with the amount of data with the AICM and HM methods. However contrary to expectations PS/SS showed a decrease in performance with 20000 bp data sets. There are several factors that can influence this behaviour, for example the path sampling chain length between the prior and the posterior, the number of sample steps and other PS/SS parameters that would need to be adjusted to a particular data set size. PS/SS are relatively new methods in phylogenetics and so far there are only a few studies investigating the influence of these parameters, generally dealing with smaller data sets and number of topologies (Lartillot & Philippe, 2006; Xie et al., 2011; Baele et al., 2013). The computational demand to investigate the possible causes of this behaviour is high and at the moment goes beyond the scope of this study. However, further research is needed especially as the genomic area will allow for the analysis of increasingly larger DNA sequence data sets.

We did not test how consistent MLL and AICM estimations are among independent MCMC runs. However Beale *et al* (2012) found that PS and SS produce consistent estimates among MCMC runs more often than the other methods. Thus, considering our results in light of previous studies (Lartillot & Philippe, 2006; Xie et al., 2011; Baele et al., 2012), we suggest that applying PS and SS would produce more reliable results than HM and AICM. However, independent of the method of hypothesis comparison used, it is always advisable to rely only on inferences with moderate to strong support.

The prior-based approach proved effective in discriminating between competing hypotheses when applied to empirical data (data set by Zhang et al., 2008). The hypotheses compared reflected scenarios well apart in time and relied on a relatively large data set and a robust phylogeny. Researchers applying this approach should meet these conditions because divergence time and tree topology are estimated at the same time with BEAST, thus changes in topology affect divergence times and vice versa (Heled & Drummond, 2011). Furthermore, as recently demonstrated, the effect of the rate priors could also affect the estimation of divergence times and should be investigated in future studies (Reis, Zhu & Yang, 2014).

In the simulated phylogenies we used a relatively high ratio of constrained/no-constrained nodes (five nodes per hypothesis comparison, plus up to three additional calibrated nodes out of 24; see input files in S2). It will be necessary to investigate if reducing the number of constrained nodes could lead to a decrease in the strength of inferences, and if an increase will improve the accuracy of the divergence estimation and thus benefit hypothesis selection. We already observed that constraining 7/40 nodes in the empirical data set analyses led to discrepancies among hypothesis selection methods. This additionally suggests that the direct comparison between these simulated and empirical data analyses should be taken with caution.

We executed unconstrained analyses that need to be run only one time to compare several hypotheses simultaneously. Most of the times, Uc was slightly outperformed by PS/SS. However the unconstrained MCMC runs reached convergence less often than the prior-based approach runs. Thus, there might not be a computational benefit in running one very long MCMC instead of several shorter parallel runs reflecting competing hypotheses. Another potential problem with just running a single run and counting the visits to each hypothesis, as we did in the unconstrained analyses, is that if the hypotheses are really disjoint, it will be necessary to throw away MCMC iterations for the times outside the hypotheses. If the hypotheses were overlapping it would be necessary to correct for this when estimating a time that could belong to different hypotheses which is an extra challenge.

The development of new methods for model selection, and future research on their performance, will add confidence to inferences led by hypothesis comparison. This could have implications for biogeographical and phylogeographical studies where robust methods for historical inferences are still lacking. Depending on the location of the nodes of interest, the approach here evaluated could also be applied in cases where not only divergence between two taxa, but instead a diversification event is suspected. At this scale, it could complement the traditional method of testing the hypothesis of shifts or heterogeneity in diversification rates against the null hypothesis of constant rates through time and among lineages (Pybus & Harvey, 2000; Chan & Moore, 2002; Ricklefs, 2007; Moore & Donoghue, 2009; Steeman et al., 2009; Silvestro, Schnitzler & Zizka, 2011). It would also allow testing the association of such shifts with climate or geological change (Hines, 2008; Schuettpelz & Pryer, 2009).

## Acknowledgements

Thanks to P. Zhang for kindly providing the Salamandridae data and the input file for BEAST. Thanks to M. Forrest for help with scripting and facilitating access to the Frankfurt Cloud (Goethe University-Deutsche Bank), and for his and C. Weiland’s assistance to access and use the BiK-F computer cluster. EZ thanks P. Jansson for providing comments.

## Supplementary Information

Table S1. Characteristics of simulation treatments.

Table S2. Properties of competing hypotheses (see also Figures 1 and 2).

File S1. Tree topologies used to generate sequence data.

File S2. xml files used as input in simulations. Available from: https://drive.google.com/open?id=0B7P6iuJv3fpiczBrQ3FDcFRGc1E

